# Massively parallel profiling and predictive modeling of the outcomes of CRISPR/Cas9-mediated double-strand break repair

**DOI:** 10.1101/481069

**Authors:** Wei Chen, Aaron McKenna, Jacob Schreiber, Yi Yin, Vikram Agarwal, William Stafford Noble, Jay Shendure

## Abstract

Non-homologous end-joining (NHEJ) plays an important role in double-strand break (DSB) repair of DNA. Recent studies have shown that the error patterns of NHEJ are strongly biased by sequence context, but these studies were based on relatively few templates. To investigate this more thoroughly, we systematically profiled ∼1.16 million independent mutational events resulting from CRISPR/Cas9-mediated cleavage and NHEJ-mediated DSB repair of 6,872 synthetic target sequences, introduced into a human cell line via lentiviral infection. We find that: 1) insertions are dominated by 1 bp events templated by sequence immediately upstream of the cleavage site, 2) deletions are predominantly associated with microhomology, and 3) targets exhibit variable but reproducible diversity with respect to the number and relative frequency of the mutational outcomes to which they give rise. From these data, we trained a model that uses local sequence context to predict the distribution of mutational outcomes. Exploiting the bias of NHEJ outcomes towards microhomology mediated events, we demonstrate the programming of deletion patterns by introducing microhomology to specific locations in the vicinity of the DSB site. We anticipate that our results will inform investigations of DSB repair mechanisms as well as the design of CRISPR/Cas9 experiments for diverse applications including genome-wide screens, gene therapy, lineage tracing and molecular recording.

## Introduction

Genome engineering tools (zinc finger nucleases, TALENs and CRISPR/Cas9) are typically used by directing endonuclease activity to a specific location in a genome, thereby introducing a double-strand break (DSB) in a directed fashion. In mammalian cells, such DSBs are primarily repaired by one of two pathways -- homology directed repair (HDR) and classical non-homologous end joining (c-NHEJ) (Hsu et al. 2014; Lieber 2010). HDR uses homologous template sequences to repair the DSB, potentially introducing programmed edits via the repair template. In contrast, c-NHEJ directly rejoins the broken ends, often perfectly but occasionally introducing errors, typically in the form of short insertions or deletions (indels) (Bétermier et al. 2014). In addition to HDR and cNHEJ, there is evidence for an alternative NHEJ pathway (alt-NHEJ), also termed microhomology mediated end joining (MMEJ), wherein short, homologous sequences in the vicinity of the DSB are used to align the broken ends prior to joining, resulting in deletions or potentially more complex events (Sfeir & Symington 2015). Below, we use ‘NHEJ’ to refer to both c-NHEJ and MMEJ/alt-NHEJ, *i.e.* template-free editing.

In recent years, CRISPR/Cas9 has emerged as a particularly versatile tool for genome editing. For many if not most applications of CRISPR/Cas9-mediated genome engineering, it is used in conjunction with the cell’s endogenous NHEJ machinery to introduce short indels in a targeted fashion (Mali et al. 2013; Cong et al. 2013; Jinek et al. 2013), *e.g.* to disrupt the function of genes or regulatory elements (Wang et al. 2015; Gasperini et al. 2017; Klein et al. 2018) or to introduce irreversible changes that record cell lineage or molecular events (McKenna et al. 2016; Frieda et al. 2017; Kalhor et al. 2018). However, despite NHEJ’s central importance to this transformative tool, our understanding of the processes that determine the rate and patterns of NHEJ-mediated errors remains incomplete.

Recent studies have demonstrated that the error outcomes of NHEJ are strongly dependent on sequence context (van Overbeek et al. 2016; Aubrey et al. 2015). Other studies show that the characteristics of the broken ends (blunt or staggered end; length of any overhang) also affect end-joining patterns both *in vitro* (Chang et al. 2016) and *in vivo* (Chang et al. 2016; Bothmer et al. 2017). However, a systematic profiling of the sequence determinants of NHEJ repair patterns has yet to be undertaken.

Here we profiled ∼1.16 million mutational events resulting from CRISPR/Cas9-mediated cleavage and NHEJ-mediated DSB repair of 6,872 synthetic target sequences. From the resulting data, we identify the primary features of sequences adjacent to the sites of DSBs that shape the distribution and relative frequency of NHEJ-mediated mutational outcomes, *e.g.* nucleotide content and microhomology. We furthermore exploit microhomology to demonstrate the “programming” of deletion patterns. Finally, we develop a statistical model to accurately predict the error patterns that result from CRISPR/Cas9-mediated cleavage of an arbitrary sequence.

## Results

### Development of a massively parallel strategy to profile NHEJ-mediated genome edits

Toward a comprehensive understanding of the sequence determinants of NHEJ-mediated error patterns, we developed a strategy that would allow us to efficiently profile a large number of repair events from each of a large number of sequence contexts (**Figure 1**). In brief, we designed 70,000 essentially random targets for CRISPR/Cas9 single guide RNAs (sgRNAs) and used array-based oligonucleotide synthesis to encode these targets in *cis* with their corresponding sgRNAs, separated by only 20 bp. We then amplified and cloned these molecules to a lentiviral vector. In our initial experiments, the complexity of the resulting library of synthetic targets and their cognate sgRNAs was such that we obtained relatively few edited templates per target. Therefore, we re-cloned the library under bottlenecking conditions, reducing its complexity to 12,917 targets. We then proceeded with viral packaging and transduction, in triplicate, of a monoclonal HEK293T cell line that stably expresses Cas9 (multiplicity of infection of ∼4-8). As such, within any given cell, only one or a few sgRNAs are expressed, and each one directs Cas9-mediated DSBs to a target located immediately adjacent to it. After five days to allow for the introduction of NHEJ-mediated errors at these targets, cells were harvested and genomic DNA isolated. We then PCR amplified the region comprising the targets and corresponding sgRNAs, using unique molecular identifiers (UMIs) appended during the first extension cycle to distinguish whether identical edits were derived from the same cell or different cells.

**Figure 1.**
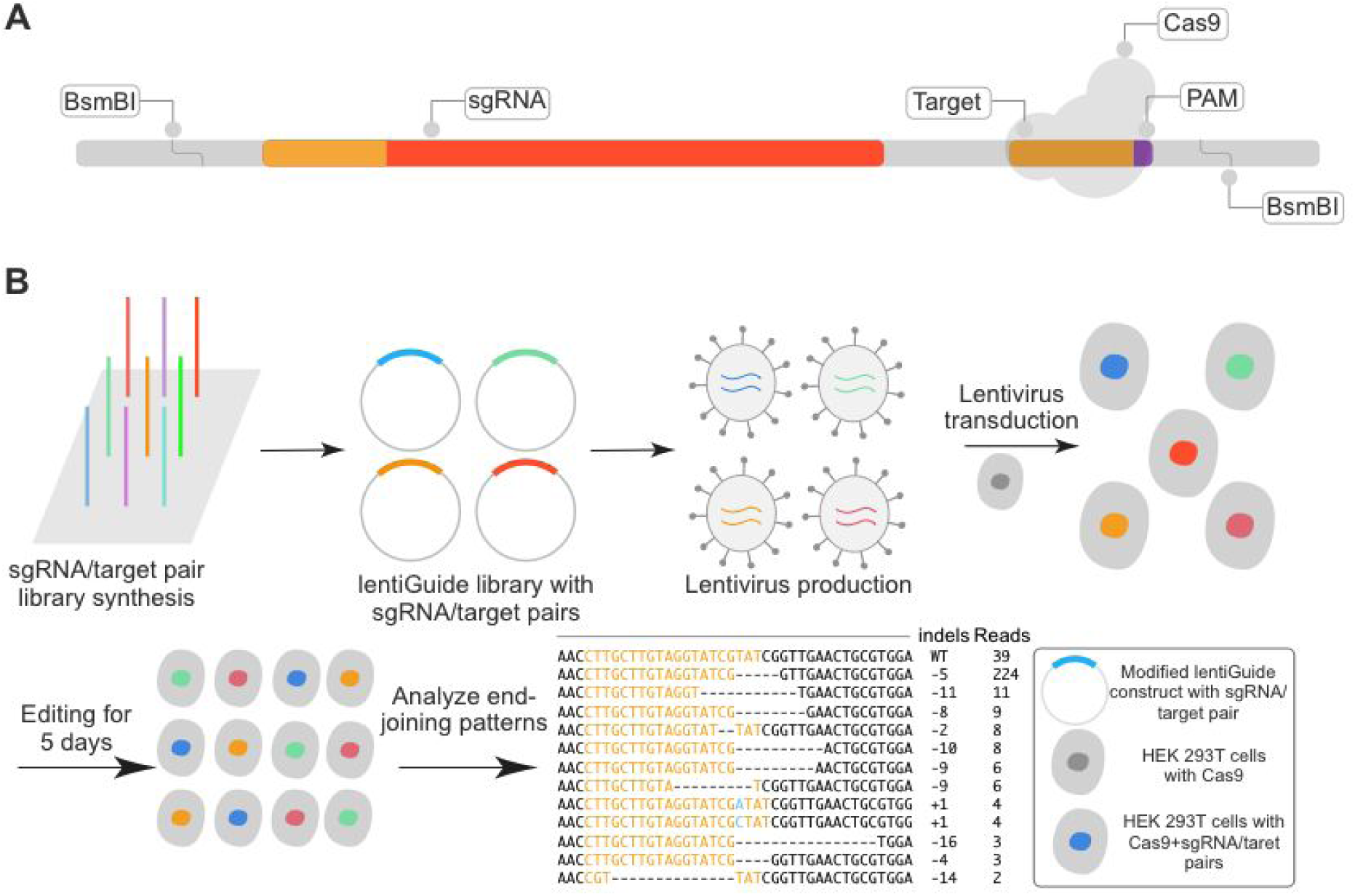
An assay for massively parallel profiling of the outcomes of CRISPR/Cas9-mediated double-stranded DNA break repair. **(A)** Schematic of library of 200 bp oligonucleotides encoding sgRNAs targeting a large number of designed 20 bp spacers, with their matched target sequence encoded *in cis*. In our primary experiment, 70,000 targets of random sequence were designed and cloned. **(B)** After array-based synthesis and PCR amplification of the library, BsmBI restriction sites at either end were used for cloning into a modified lentiviral construct. The library was bottlenecked to 12,286 targets to facilitate greater coverage of independent NHEJ-mediated events corresponding to each target. Monoclonal HEK293T cells expressing Cas9 were transduced with packaged lentivirus. Cells were harvested at 5 days after transduction, and a region including both the spacer and the target was PCR amplified from genomic DNA for high-throughput sequencing. The sequences of mutated targets were aligned to their corresponding unmutated reference (assigned based on the spacer sequence).

Summing across the three replicates, we sequenced PCR amplicons to a depth of ∼148 million reads, which were reduced to ∼1.19 million reads after collapsing on the basis of identical sequences and UMIs, and filtering of reads with evidence of lentivirus-mediated template switching (Sack et al. 2016; Hill et al. 2018) or other unexpected sequences (*e.g.* synthesis or PCR errors). After further filtering of poorly represented targets (those represented by fewer than 10 UMIs), our dataset consisted of ∼1.16 million UMIs corresponding to 6,872 unique targets. On average, each target was represented by 168 UMIs and 24 alleles (where ‘allele’ refers to a unique post-editing sequence of a given target). Each allele was aligned to its original sequence, known because the corresponding spacer sequence is part of the same amplicon, using the Needleman-Wunsch algorithm (Needleman & Wunsch 1970). Alleles were categorized as wild-type (*i.e.* unedited), a deletion, or an insertion.

Overall, targets were highly edited, with only 9.8% of UMIs corresponding to the wild-type allele. Of UMIs containing detectable mutations, 63.6% were deletions and 31.5% were insertions (**Figure 2A**). The remainder (4.9%) contained some combination of substitutions, insertions and deletions, and are excluded from all of our subsequent analyses. Deletions were dominated by small events; only 1.5% were >25 bp, although we note that deletions >150 bp are not captured by our assay (Gasperini et al. 2017; Kosicki et al. 2018). The vast majority of insertion events were of a single base pair.

**Figure 2.**
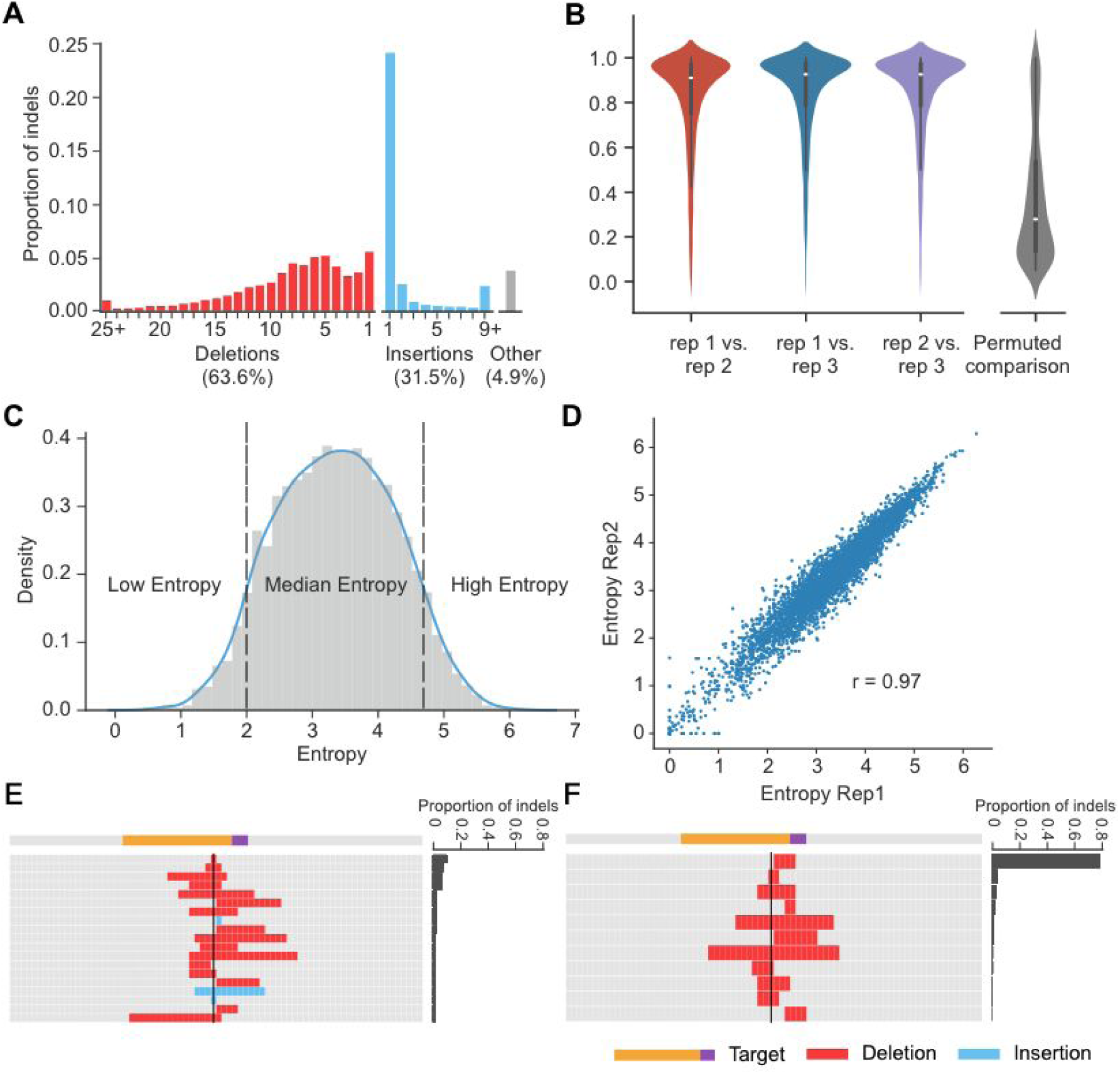
Mutation patterns resulting from DSB repair vary greatly between targets, but are highly reproducible for individual targets. **(A)** Overview of indel profiles. Of all detectably mutated targets, 63.6% were small deletions (red) and 31.5% were insertions (blue). The remainder (4.9%) contained some combination of substitutions, insertions and deletions, and are excluded from all subsequent analyses. **(B)** End-joining patterns were highly reproducible for the same target between replicates. Left: violin plot of distribution of correlation coefficients for pairwise comparison of individual targets between replicates. Right: Permuting the allele counts for each target in one replicate and repeating the pairwise comparison greatly reduces the observed correlations. **(C)** Entropy quantifies the diversity of NHEJ outcomes from individual targets. Targets were separated into low, medium and high entropy classes. **(D)** Estimated entropy for individual targets was highly reproducible between replicates (rep1 vs. rep2 shown). **(E, F)** Example of targets with high and low entropy. High entropy targets had diverse outcomes at appreciable frequencies (E) while low entropy targets were dominated by a single outcome (F).

### Repair patterns are reproducible but exhibit highly variable ‘entropy’ between targets

We sought to examine whether repair patterns for any given target were reproducible, as previously shown for a more limited set of templates (van Overbeek et al. 2016). For each target, we calculated the frequency of each non-wild-type allele. For any given target, the distribution of frequencies for its alleles were highly reproducible in pairwise comparisons of the three replicates (median Pearson’s r = 0.91, 0.93, 0.93, **Figure 2B**, left). Meanwhile, if we permute the alleles in one replicate on a target-by-target basis and repeat the pairwise comparison, these correlations are greatly reduced (median Pearson’s r = 0.20, **Figure 2B**, right).

Confirming the observations of (van Overbeek et al. 2016), the diversity of mutations strongly varied from target to target. We calculated the Shannon entropy of mutational outcomes for any given target as -∑ p_i_*log(p_i_), where p_i_ is the frequency of i^th^ indel of that target (**Figure 2C**). Entropy values for any given target were highly reproducible between replicates (**Figure 2D**) and only modestly correlated with sampling depth (**Figure S1**). Of note, some targets consistently exhibited particularly diverse mutational outcomes consequent to NHEJ -- that is, high entropy (*e.g.* **Figure 2E**, where the most frequently observed mutation occurs in only 10.1% of mutated templates). Other targets were strongly biased towards a more limited set of mutational outcomes -- that is, low entropy (*e.g.* **Figure 2F**, where the most frequently observed mutation occurs in 80.4% of mutated templates).

### Sequence context at the DSB site predicts the frequency of insertions

We next sought to investigate the determinants of insertions at the DSB, which are dominated by 1 bp events (**Figure 3A**). 84% of 1 bp insertions were predicted (and presumably templated) by the nucleotide immediately upstream of the cleavage site (*i.e.* the 17th nucleotide in target sequence; **Figure 3B**; **Figure S2**) Although it might have been expected that NHEJ-mediated repair would be symmetric with respect to the site of a DSB, we do not observe templating from the immediately downstream (18th) nucleotide (**Figure 3B**). Similarly, of 2 bp insertions, a substantially greater than expected proportion (41%) were templated by the sequence immediately upstream of the DSB (*i.e.* inserted sequence identical to the 16th and 17th nucleotides of the target sequence; **Figure 3C**). The asymmetric templating of NHEJ-mediated insertions was also described in two other recent studies based on data from yeast and mice (Lemos et al. 2018; Kalhor et al. 2018).

**Figure 3.**
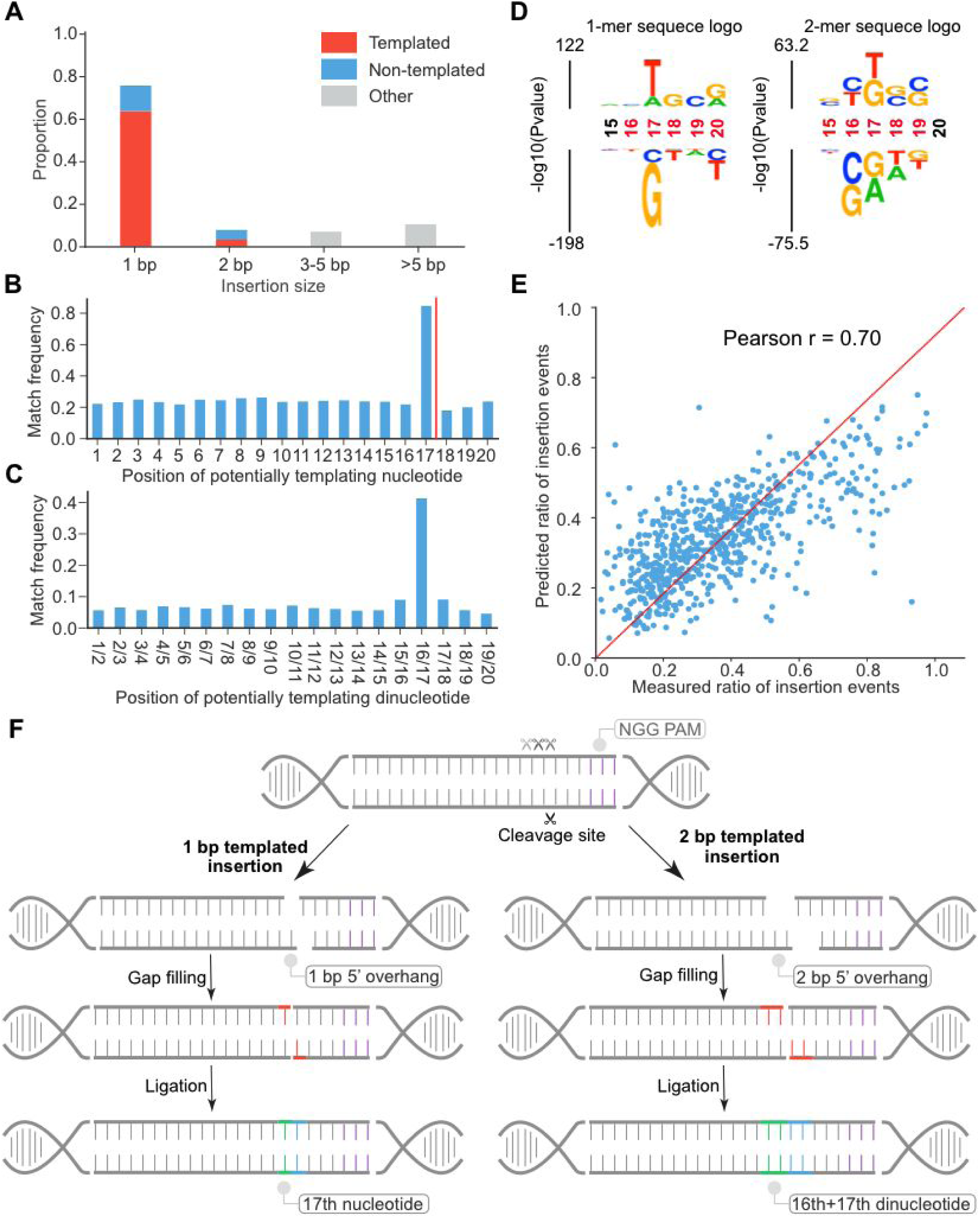
A model for asymmetric templating of NHEJ-mediated insertion events at sites of CRISPR/Cas9-mediated DSBs. **(A)** 75.3% of the insertions were 1 bp. Of 1 bp insertions, 85% appear to be templated. **(B, C)** Histogram of the number of 1 bp (B) or 2 bp insertion events where the inserted base or dinucleotide is identical to the base at a specific position in the target. The canonical DSB site is between 17^th^ and 18^th^ of the target sequence (red line). The result suggests many 1 bp insertions are templated by the nucleotide at the 17^th^ position but not the 18^th^ position (B) and many 2 bp insertions are templated by dinucleotide at the 16^th^ and 17^th^ positions (C). **(D)** The immediate sequence context surrounding the DSB strongly biases the proportion of NHEJ-mediated outcomes that result in insertions vs. deletions. The 1-mer sequence logo (left) shows that the presence of a ‘T’ and ‘A’ at the 17^th^ position increased the ratio of insertions. The 2-mer sequence logo (right) shows that the presence of a ‘TG’ dinucleotide at the 17^th^/18th position increased the ratio of insertions, while a ‘CG’ dinucleotide at the 16^th^/17th position, or a ‘GA’ dinucleotide at the 17^th^/18th position, decreased the ratio of insertions. Significant positions are colored in red. **(E)** A regression model using the nucleotide content of a 6 bp window centered on the DSB site predicted the ratio of insertion-to-deletion events. **(F)** A model for how insertions at CRISPR/Cas9-mediated DSBs are asymmetrically biased by local sequence context. Local sequence context biases the pattern of cleavage of the non-complementary strand to the sgRNA, resulting in different frequencies of blunt vs. 5’ overhangs for different targets. This in turn biases the ratio of insertions vs. deletions, as 5’ overhangs are preferably repaired by gap-filling (red) and ligation, resulting in the observed preponderance of 1 bp or 2 bp templated insertions (green/blue).

Because the ratio of insertions to deletions varied from target to target, we used kpLogo (Wu & Bartel 2017) to examine what local sequence features might shape this. We find that the presence of a T or A at the 17th bp of the target was associated with insertion events, while a G or C at this position was associated with deletion events (**Figure 3D**, left). Additional analyses showed a ‘TG’ dinucleotide flanking the cleavage site to be the most highly biased toward insertion (57% of events with that context are insertions), while a ‘GA’ dinucleotide flanking the cleavage site was the most highly biased towards deletion (17% of events with that context are insertions) (**Figure 3D**, right).

We split 2,680 targets associated with both insertion and deletion outcomes into training (n = 2,000) and test (n = 680) sets, and trained a linear regression model to predict the proportion of insertion events based on position-specific content of the hexamer centered on the DSB (single and dinucleotide k-mers; 104 binary features; **Figure 3E**). The model performs reasonably well (Pearson’s r = 0.70).

Overall, these analyses confirm that local sequence around the DSB site plays an important role in shaping the outcome(s) of NHEJ-mediated errors. In particular, the asymmetry implied by the high rate of identity between 1-2 bp insertions and the nucleotides immediately upstream to the DSB, but not the nucleotides immediately downstream to the DSB (**Figure 3B-C**), suggests that not all CRISPR/Cas9-mediated cleavages are blunt-ended. Indeed, *in vitro* studies have shown that the non-complementary strand of the target can sometimes be cleaved by Cas9 at multiple sites upstream of the −3 bp position relative to the protospacer adjacent motif (PAM), while the complementary strand is cut only at that site, instances which would result in a 5’ overhang (Jinek et al. 2012; Stephenson et al. 2018). The preponderance of 1 bp insertions templated by the 17th rather than 18th base could be explained by fill-in of this overhang followed by blunt-ended ligation (and similarly for the preponderance of 2 bp insertions that are templated by the 16th and 17th bases, rather than the 18th and 19th bases).

To summarize, we propose a model (**Figure 3F**) where 1) some proportion of cleavages of the non-complementary strand by Cas9 occur upstream of the −3 bp PAM cleavage site, while cleavage of the complementary strand always occurs between the 17th and 18th positions, resulting in a 5’ overhang; 2) 5’ overhangs are preferably repaired by gap-filling and ligation, resulting in the observed bias towards templating by the bases immediately upstream rather than downstream of the DSB; 3) local sequence context biases the pattern of cleavage on the non-complementary strand, resulting in different frequencies of blunt vs. 5’ overhangs for different targets, which in turn biases the ratio of insertions vs. deletions. A similar model was recently proposed by (Lemos et al. 2018) based on asymmetric templating of NHEJ-mediated insertions observed in yeast.

### Extensive use of microhomology in NHEJ-mediated deletions

We next examined patterns of deletion. Microhomology (MH) refers to the use of short regions of identical sequence (1 to 16 bp) that can mediate the alignment of broken ends (**Figure 4A**) and is relevant to both c-NHEJ and alt-NHEJ/MMEJ (Deriano & Roth 2013; Pannunzio et al. 2014; Sfeir & Symington 2015; Conlin et al. 2017). Here, a deletion event is considered to be MH-mediated if the sequence at the 3’ of a rejoined end is identical to the 3’ end of the deleted sequence, and the size of the MH tract refers to the length of that identical sequence. By that definition, we found that over 75% of deletion events in our dataset are MH-mediated. The length of MH tracts ranged from 1-10 bp. Nearly all (94.6%) MH-mediated events involved relatively short tracts of microhomology, *i.e.* 1-4 bp. Longer MH tracts were observed more rarely (**Figure S3A**), probably simply due to the relative paucity of opportunities in our set of random target sequences.

**Figure 4.**
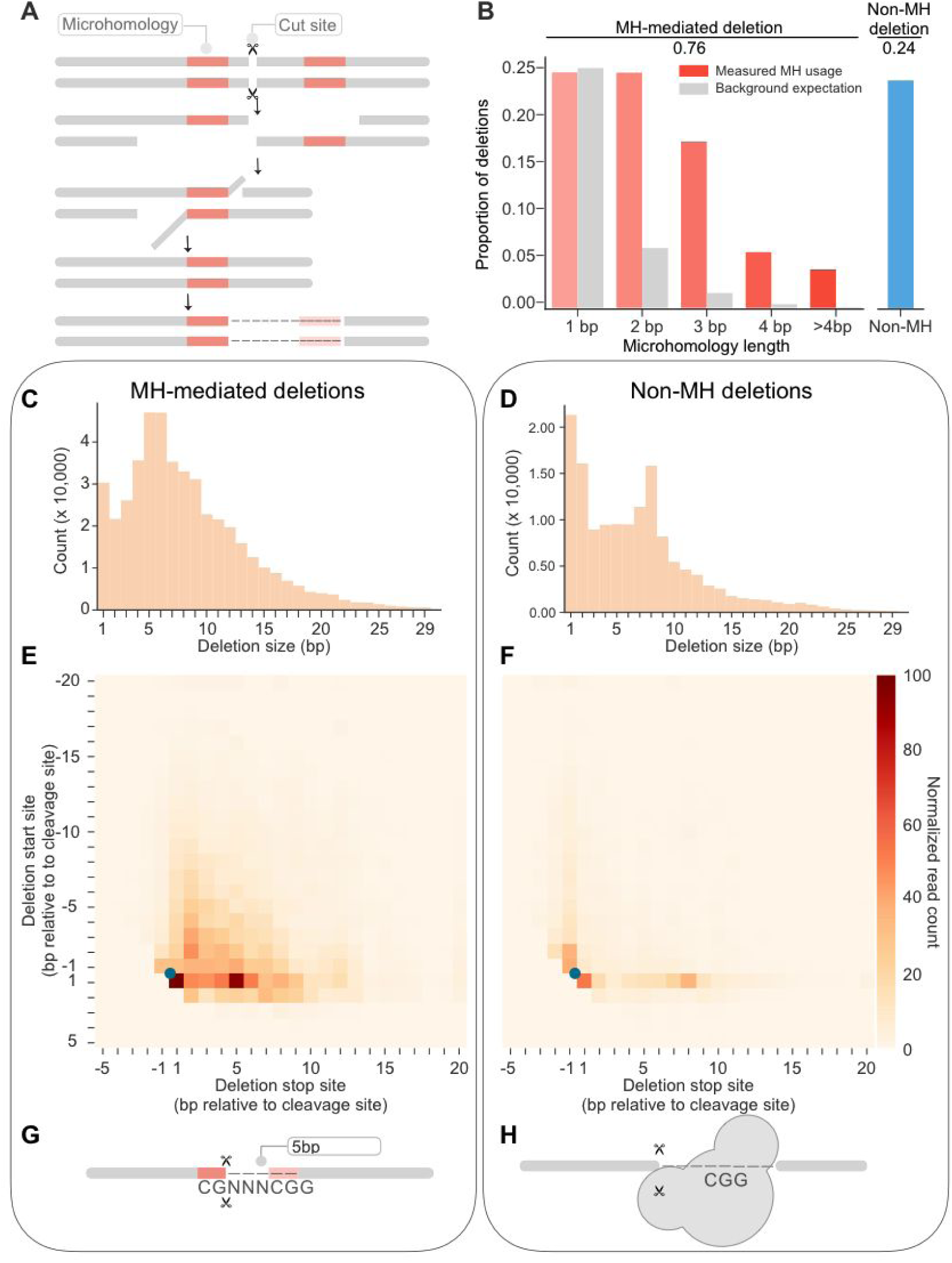
Extensive use of microhomology in NHEJ-mediated deletion events. **(A)** Schematic of microhomology (MH) usage in end-joining repair. Tracts of MH (red) in the vicinity of the DSB are used to align the broken ends. The unannealed overhang is cleaved by endonuclease and the gap filled by polymerase. Here, a deletion event is defined as MH-mediated deletion if the sequence at the 3’ of a rejoined end (red, left) is identical to the 3’ end of the deleted sequence (red, right). The size of the MH tract refers to the length of that identical sequence. **(B)** Length distribution of MH tracts in observed MH-mediated events. With the exception of 1 bp deletions, all MH tract lengths occured at substantially greater than expected frequencies. **(C, D)** Distribution of deletion sizes of MH-mediated (C) and non-MH (D) events. **(E, F)** Heatmap of showing frequency of start/stop sites of MH-mediated (E) and non-MH (F) deletion events. The Y and X axes correspond to the start and stop sites of deletion events, respectively, with positions shown relative to the canonical DSB site (blue dot). Both MH-mediated and non-MH deletions were primarily “unidirectional” relative to the DSB site, rather than spanning it. **(G)** Schematic of potential explanation for the observed excess of 5-6 bp MH-mediated deletions. PAM-like sequences near the DSB are biased towards deletion events. G: Microhomology between a G at the 17th position or a CG at the 16th/17th position with corresponding sequences in the PAM result in an excess of 5-6 bp deletions. **(H)** Schematic of potential explanation for the observed excess of 8 bp non-MH deletions. In the dsDNA-sgRNA-Cas9 complex, the region 1-8 bp downstream of the cleavage site is occupied by Cas9. The enrichment of non-MH deletions 8 bp from the cleavage site could simply correspond to the nearest position lacking Cas9 protection from endonucleases during repair.

The frequencies of tracts of various lengths consistent with MH usage were substantially higher than background expectation for all lengths except 1 bp, with that enrichment increasing as a function of tract length (**Figure 4B**). We further investigated the relevance of 1 bp MH by comparing the proportion of 1 bp deletion events in targets with identical vs. non-identical nucleotides immediately spanning the cleavage site. We observe a 3-fold greater proportion of 1 bp deletion events when those nucleotides are identical than when they are not (**Figure S3B**), suggesting that 1 bp MH may play a role in aligning, stabilizing and rejoining the broken ends.

The lengths of MH vs. non-MH mediated deletions exhibited distinct distributions (**Figure 4C-D**). In particular, the distribution of deletion sizes for MH-mediated events peaks at both 1 bp and 5-6 bp, while an equivalent distribution for non-MH-mediated deletions peaks at both 1-2 bp and 8 bp. The frequency of longer deletions exhibits an exponential decay for both MH and non-MH mediated events. To investigate this further, we jointly analyzed the frequency of start and end points for deletion events, relative to the position of the canonical cleavage site (**Figure 4E-F**). Both MH and non-MH mediated deletions exhibited a preference for “unidirectional” events, *i.e.* either the start or end point is immediately adjacent to the cleavage site, rather than the deletion spanning the cleavage site.

What explains the excess of deletion events of specific lengths? For MH-mediated events, the excess of 1 bp deletions may simply be attributable to the aforedescribed instances of identical nucleotides spanning the cleavage site (**Figure S3B**). However, the excess of 5-6 bp MH-mediated events is clearly driven by events in the downstream direction (**Figure 4E**), *i.e.* deletions between the DSB and the PAM. A potential explanation is that the predilection of PAM-like sequences near the DSB for deletion events (*i.e.* a G nucleotide at the 17th position or a CG dinucleotide at the 16th/17th position; **Figure 3D**), coupled with the consistent presence of the CGG PAM sequence at the 21st-23rd position, results in an excess of deletions mediated by CG (5 bp deletion) or G (5-6 bp deletion) microhomology (**Figure 4G**).

For non-MH-mediated events, the excess of 8 bp events might be explained by the observation that in the dsDNA-sgRNA-Cas9 complex, the region 1-8 bp downstream of the cleavage site is occupied by Cas9, even after cleavage (Jiang et al. 2016; Stephenson et al. 2018). Thus, the enrichment of non-MH deletions 8 bp from the cleavage site could simply correspond to the nearest position lacking Cas9 protection from endonucleases during repair (**Figure 4H**).

### Generating predictable mutations by programming microhomology tracts

Since MH is widely used in deletion events, we reasoned that we could program a library of targets to generate predictable mutations by introducing MH proximal to the cleavage site. With the same basic experimental scheme (**Figure 1**), we tested a library of 1,000 targets and corresponding guides containing MH tracts of three different lengths (2, 4 or 6 bp) matching the sequence immediately upstream of the expected DSB site, and positioned 6 bp downstream of the cleavage site (**Figure 5A-B**). The resulting data were processed and analyzed similarly to the previous experiment.

**Figure 5.**
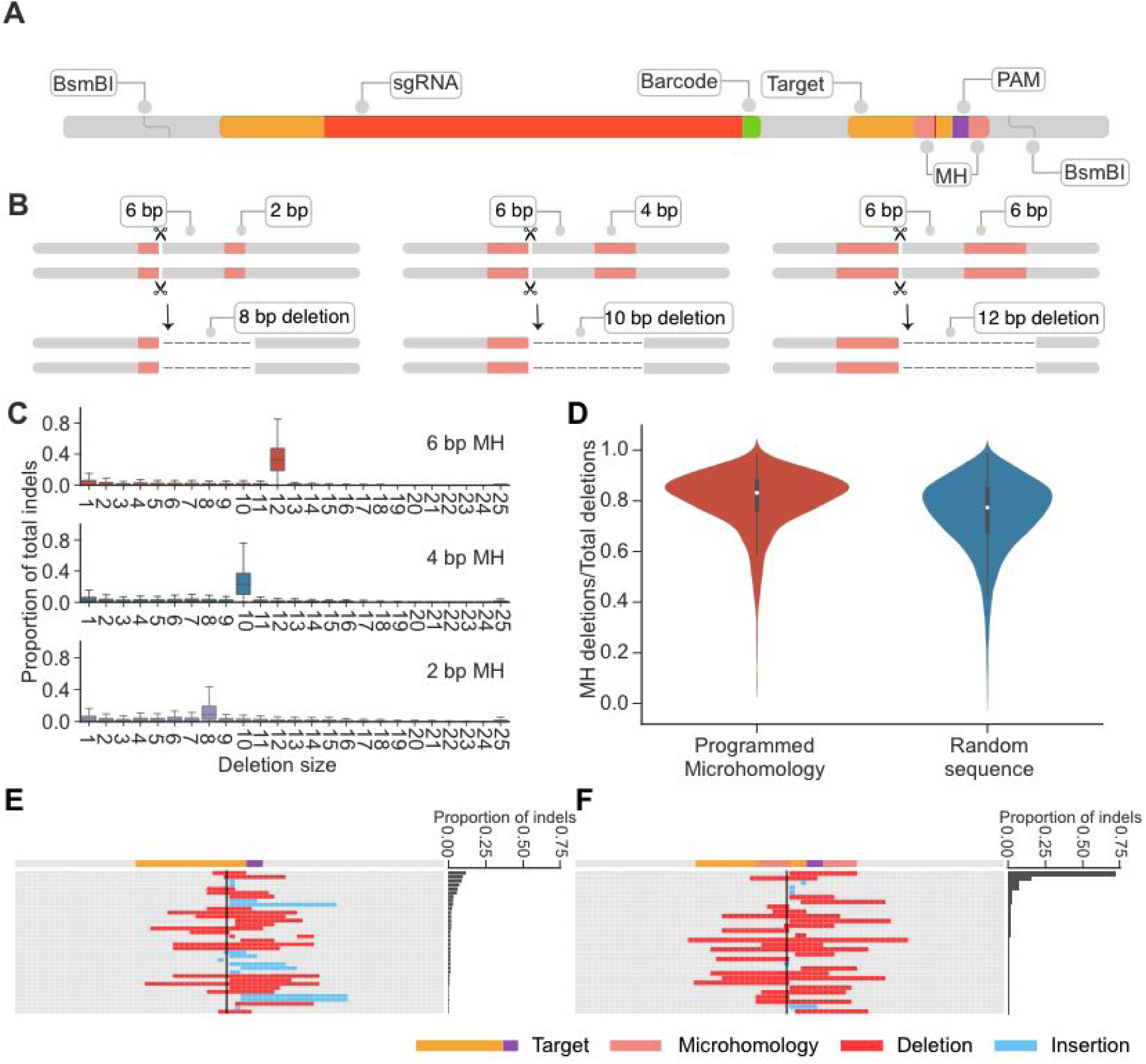
Programming microhomology tracts into targets increases predictability of repair outcomes. **(A,B)** Schematic of programmed MH tract designs and expected deletion sizes. The distance between the regions of MH (red) was consistently 6 bp, while the MH tracts were 2 bp, 4 bp or 6 bp, such that the expected deletion sizes were 8 bp, 10 bp and 12 bp, respectively. **(C)** Distribution of observed deletion sizes for targets with programmed MH tracts of various lengths. We observe a strong bias towards deletions of the expected lengths, with the proportion increasing for longer MH tracts. **(D)** MH usage in sequences with (left) or without (right) programmed MH. Despite the strong bias of towards intended deletions when MH occurred, the proportion of MH events only slightly increased from 76% to 82%. **(E,F)** Example of a sequence that shows diverse editing outcomes (E). However, when a 6 bp MH tract is introduced onto this sequence backbone, the programmed 12 bp deletion comprises nearly 75% of the editing outcomes (F).

Intentionally programming MH tracts resulted in a high proportion of events corresponding to the expected deletions (8, 10 and 12 bp deletions for 2, 4 and 6 bp MH tracts, respectively; **Figure 5B-C, Figure S4**). We also observe that the ratio of the programmed deletion increases as a function of length of the MH tract (**Figure 5C**). However, despite the greater predictability of which MH-mediated outcome would occur, the relative proportion of MH-mediated deletions increased only slightly from 76% to 82% (**Figure 5D**). Furthermore, we did not observe an excess of ‘imperfect’ MH-mediated events, *e.g.* an excess of 11 bp or 13 bp deletions in targets for which a 12 bp deletion was expected (**Figure S4**). Nonetheless, the results show how targets that would result in diverse editing outcomes can be strongly biased towards a specific outcome by the presence of MH tracts (**Figure 5E-F**).

### A machine learning model to predict editing patterns

The above results above suggest that the NHEJ-mediated repair outcomes for any given target sequence are both reproducible and dependent on sequence context. Accordingly, we next sought to train a machine learning model to predict these outcomes and their relative frequencies. We began by filtering out target sequences that were either poorly reproducible (low correlation between replicates, mainly due to low UMI counts; **Figure S5A**) or poorly edited, resulting in a dataset of ∼1 million UMIs representing 4,611 target sequences. On average, each target in this subset of the data used for modeling was represented by 215 UMIs and 24 alleles.

Because larger events are rare in our data, we focused on predicting deletion events ≤ 30 bp in length, as well as all possible 1-2 bp insertion events at the DSB. Across all targets, we identified 584 “event classes”. The vast majority of CRISPR/NHEJ-mediated indels arising from any given target sequence should fall into one of these 584 event classes. We therefore framed our machine learning task as one of predicting, for an arbitrary target sequence, the relative frequency of CRISPR/NHEJ-mediated indels falling into each of these 584 event classes. These included 563 deletions (defined solely by their start/end points), all 4 possible single nucleotide insertions, all 16 possible dinucleotide insertions, and finally, a single event class for insertions greater than 2 bp in length. Of note, the 563 deletion event classes comprise almost all of the 585 possible combinations of start/end positions, with the constraints that deletions must be less than 30 bp and overlap with the −3/+2 window around the cleavage site. The 22 potential deletions that satisfy these constraints but were not observed in the modeling dataset were mainly large deletions.

We also defined 2,962 binary features to characterize the target sequence for which repair outcomes are being predicted. These are 1) 384 binary features corresponding to one-hot encoded sequence, including 80 for single nucleotide content (4 nucleotides * 20 positions) and 304 for dinucleotide content (16 dinucleotides * 19 positions); 2) 2,578 binary features corresponding to MH tracts; specifically, for each of the possible deletion event class, we defined 5 binary features corresponding to the length of the MH tract, if any (0-4 bp * 563 deletion event classes = total 2,815 binary features, or 2,578 after excluding 237 binary features corresponding to characteristics never observed in the training data) (**Figure 6A**).

**Figure 6.**
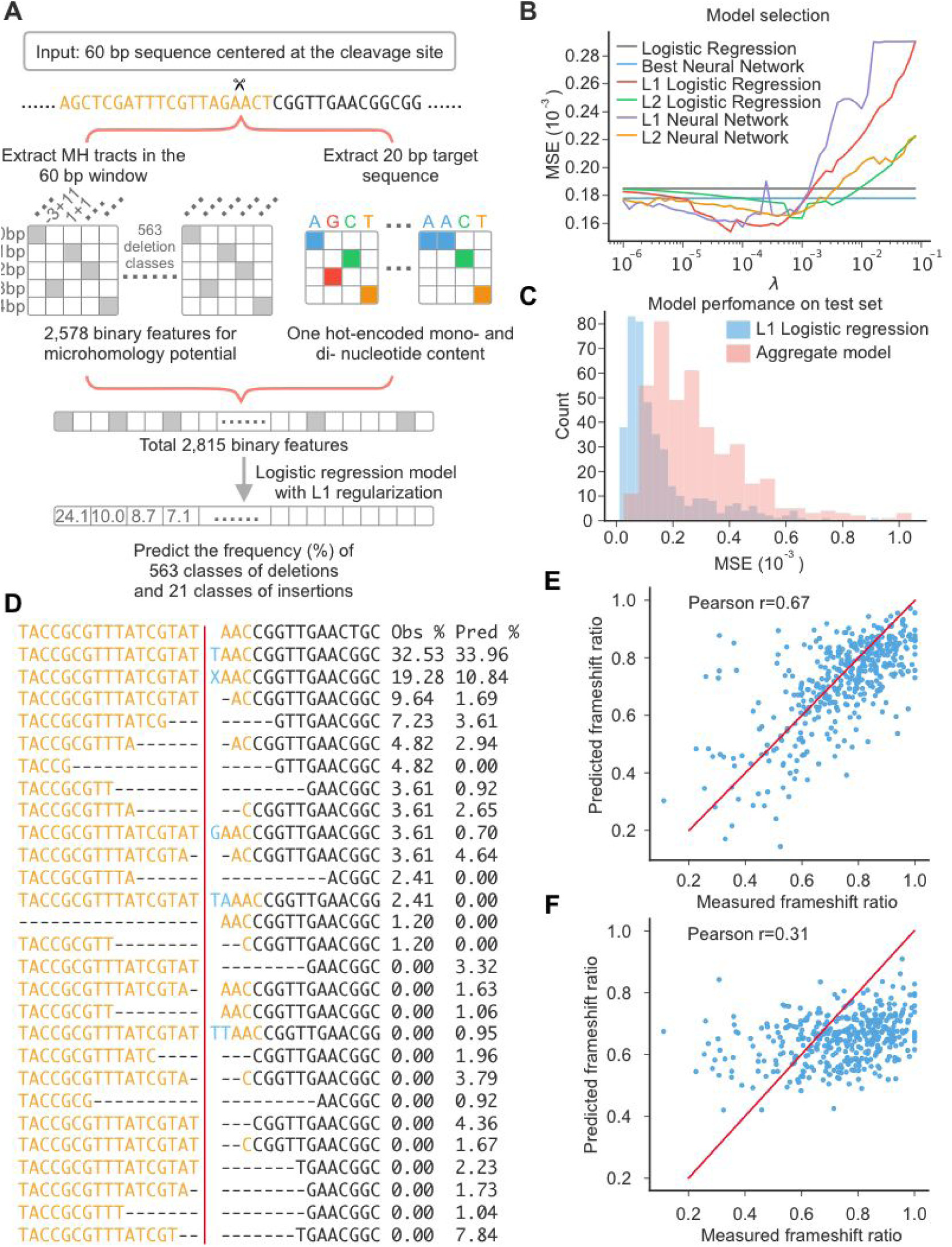
End joining patterns can be accurately predicted. **(A)** Schematic of machine learning training. A 60 bp sequence (± 30 bp around the cleavage site) is used as the input to the model. A total of 2,815 binary features -- 2,578 corresponding to MH potential and 384 to one-hot encoded mono- and di-nucleotide content of the 20 bp target -- are extracted. **(B)** Model selection. We evaluated both logistic regression and neural network models with or without regularization. A logistic regression model with L1 regularization outperformed alternative models on predicting the relative frequencies of various event classes on the validation dataset. **(C)** Performance of L1 logistic regression on the test dataset. The distribution of MSE values for the 411 test targets is shown (blue). Poorly predicted targets largely correspond to those that were poorly sampled (see Figure S5C). As a baseline to illustrate the improvement conferred by the L1 logistic regression model, we show a similar distribution for the aggregate model, in which the predicted frequencies of 584 indel classes are simply taken from the aggregate frequency at which each is observed in the training and validation datasets (red). **(D)** Example of predicted vs. observed frequencies for a specific target. The cleavage site is shown as a vertical red line. Insertions are shown in blue. The single event class corresponding to insertions >2 bp is represented as blue X. **(E-F)** The logistic regression model (E) compared favorably to Microhomology Predictor (Bae et al. 2014) (F) in predicting the ratio of frameshifting mutations for each of the 411 targets.

We split the 4,611 target sequences in our modeling dataset into subsets of 3,750 (for training), 450 (for validation) and 411 (for testing). We first evaluated different machine learning approaches, including logistic regression as well as neural networks with varying complexity (see Methods), as well as both approaches with regularization using L1 and L2 penalties. We trained each model on the training set using cross-entropy loss and evaluated performance on the validation set using the mean squared error (MSE). We found that the best performing model was a L1-regularized logistic regression (**Figure 6B, Figure S5B**), and chose to move forward with that.

Applying this model to the test set of 411 target sequences, which had been entirely held out from the training and validation steps, we compared the observed versus predicted frequencies of indels falling into various “event classes”. Observations and predictions were well matched for most targets, with a MSE of 0.00015 (**Figure 6C-D**). As a baseline, we also generated a set of predictions based simply on the aggregate frequencies of event classes in the training and validation datasets; as expected, these predictions performed more poorly (MSE of 0.00028; **Figure 6C**), confirming the improvement conferred by the model. Poorly predicted targets tended to be those with relatively shallower sampling of editing events, *i.e.* where our observed frequencies are noisier (**Figure S5C**).

Overall, the above results show that our model can accurately predict the relative frequencies of repair outcomes. As a common use of CRISPR/Cas9 in conjunction with NHEJ is to introduce frameshifting mutations, we also assessed the observed versus predicted ratios of frameshifting indels for each of the 411 targets, including observations and predictions in all event classes, and found them to be well correlated (Pearson’s r = 0.67; **Figure 6E**). This result compares favorably with the predictions of another tool that we tested on this same task for the 411 targets ((Bae et al. 2014); Pearson’s r = 0.31, **Figure 6F**).

## Discussion

In summary, we developed an assay to systematically profile the diversity and relative frequencies of mutational events resulting from CRISPR/Cas9-mediated cleavage and NHEJ-mediated DSB repair of thousands of synthetic sequences. In applying this assay and analyzing the editing outcomes associated with 6,872 random target sequences, we confirm that CRISPR/NHEJ-mediated repair outcomes for any given target sequence are reproducible, predictable, and largely shaped by the sequence context around the cleavage site (van Overbeek et al. 2016; Aubrey et al. 2015).

Our results also provide further insights into NHEJ-mediated repair of CRISPR/Cas9-mediated DSBs in human cell lines. <u>First</u>, we observe that insertion events are dominated by 1-2 bp insertions templated by the sequence immediately upstream of the cleavage site. Together with *in vitro* data from the literature (Stephenson et al. 2018; Jinek et al. 2012), the data supports a model in which the sequence context around the DSB biases the extent to which cleavages are blunt-ended vs. include a 1-2 bp 5’ overhang. Such 5’ overhangs are repaired by gap-filling and ligation, resulting in asymmetrically templated 1-2 bp insertions. <u>Second</u>, we observe extensive usage of 1-4 bp microhomology in mediating deletion events, and furthermore show that repair outcomes can be strongly biased towards predictable outcomes by intentionally introducing MH tracts at specific distances from the DSB. Notably, however, the introduction of MH tracts did not substantially increase the proportion of MH-mediated events. Third, both MH and non-MH-mediated deletions were overwhelmingly unidirectional (*i.e.* extending either upstream or downstream from the DSB, rather than spanning it).

From these data, we trained an L1 regularized logistic regression model that accurately predicts the relative frequency of repair outcomes for any given target sequence. While this manuscript was in preparation, several similar studies appeared as publications (Shen et al. 2018) or preprints (Allen et al. 2018; Chakrabarti et al. 2018). Chakrabarti et al. profiled the mutational outcomes associated with 1,492 target sequences from the human genome, while Allen et al. and Shen et al. designed and profiled 6,568 and 1,872 synthetic target sequences, respectively (Allen et al. 2018; Chakrabarti et al. 2018; Shen et al. 2018). These studies similarly identified sequence context surrounding the DSB, including MH, as a major determinant of repair outcomes, and also trained models to predict the frequency of the most common allele (Chakrabarti et al.) or the relative frequencies of all repair outcomes (Allen et al., Shen et al.) with comparable performance to that of our model.

In addition to insights into into NHEJ-mediated repair of CRISPR/Cas9-mediated DSBs, our study also provides a new tool for sgRNA design for diverse goals. First, for applications relying on gene knockout, *e.g.* CRISPR screens, the model’s accuracy in predicting which sgRNAs/targets are likely to result in a high proportion of frameshifting indels will be useful. Second, for applications focused on mutation correction (*e.g.* using CRISPR/NHEJ to correct pathogenic mutations), the model may be useful for identifying sgRNAs/targets for which the desired outcome is predicted to occur at a high or sufficient frequency. Third, we and others have recently repurposed CRISPR/Cas9 as a tool for lineage tracing and/or molecular recording (McKenna et al. 2016; Frieda et al. 2017; Kalhor et al. 2018). For some goals (*e.g.* lineage tracing), the identification of “high entropy” targets may critically enable the diversity necessary to uniquely label millions or billions of cells. For other goals (*e.g.* molecular recording), the design of “low entropy” targets may facilitate predictable sequential editing. More generally, a deeper understanding of CRISPR/NHEJ-mediated mutations will strengthen our ability to precisely orchestrate not only the locations but also the outcomes of genome editing.

## Materials and Methods

### sgRNA and target pair library design

To generate a library of CRISPR/Cas9 targets that could safely be characterized within human cells, we evaluated ∼1 million random 20mer crRNA sequences, scoring them against the human genome (version hg19) for off-target effects using FlashFry (McKenna & Shendure 2018). We excluded guides with less than 2 bp mismatch (including an exact match) to any target within the human genome or those with a off-target score less than 90 (Hsu et al. 2013), resulting in a modest bias towards targets containing CpG dinucleotides (**Figure S6**), and then selected a final library of 70,000 top scoring guides for synthesis. The resulting sgRNA sequence and their corresponding targets were separated by a common 20 bp spacer sequence and ordered as an Agilent SureGuide Unamplified Custom CRISPR Library array (**Figure 1A**).

To analyze the potential impact of programmed microhomology, we selected a subset of 1,000 sgRNA-target pairs from the library above and introduced microhomology with different lengths (2 bp, 4 bp, and 6 bp) matching the last 2, 4, and 6 nucleotides upstream of the cleavage site. Each design was assigned a 4 bp barcode, indicating its programmed microhomology pattern (**Figure 5A-B**). This library of microhomology sequences was ordered as an oligo pool from Twist Biosciences.

### Library Cloning

The lentiGuide-Puro (Addgene #52963) vector was modified with two rounds of PCR to remove the existing tracrRNA and filler sequence (primer P1, P2), and to incorporate two BsmBI restriction site for integration of sgRNA-target pairs (primer P3, P4). The modified vector was digested with BsmBI (NEB, Buffer 3.1) at 55°C for 3h and gel purified with Monarch DNA Gel Extraction Kit (NEB). This digested and purified vector was used for all downstream cloning.

Oligos with sgRNA-target pairs from Agilent or Twist Bioscience were both resuspended to 10ng/μl. The oligo pool was PCR amplified using KAPA Biosystems HiFi HotStart ReadyMix 2x using primers P5 and P6 and cleaned with the DNA Clean& Concentrator kit (Zymo Research). The purified PCR product was then digested with BsmBI (NEB, buffer 3.1) at 55°C for 1h to generate compatible sticky ends matching the modified lentiGuide-Puro above, and subsequently cleaned with DNA Clean& Concentrator (Zymo Research). Digested vector and insert were ligated with T4 ligase (NEB) with a molar ratio of 1:3. Ligation products were transformed in to Stable Competent *E.coli* (NEB C3040H). Transformed cells were cultured at 30°C overnight and plasmid DNA was prepared using a ZymoPURE II Plasmid Kit.

### Cell Culture and lentivirus transduction

We generated a mono-clonal 293T cell line expressing Cas9 by transduction of Cas9-blast lentivirus particles (Addgene plasmid #52962). Cells were cultured in DMEM High glucose (GIBCO) supplemented with 10% Fetal Bovine Serum (Rocky Mountain Biologicals) and 1% penicillin-streptomycin (GIBCO) and grown with 5% CO^2^ at 37°C.

All lentivirus libraries were produced by the Fred Hutchinson Cooperative Center for Excellence in Hematology Vector Production core facility. HEK293T cells were transduced and media was changed to virus free media at 24 hours post-transduction. Cells were passed every 48h with a split ratio of 1:6. Cells were harvested at day 5 after transduction.

### Sequencing Library Generation

Genomic DNA was extracted with DNeasy Blood & Tissue Kit (Qiagen) following the manufacturer’s protocol. 15 bp unique molecular identifiers (UMIs) were added by one initial round of linear PCR using a primer containing a 5′ sequencing adaptor (P7). For each reaction we used 250ng of genomic DNA, 0.2μl 100mM primer and 25 μl HiFi HotStart ReadyMix 2x (KAPA Biosystems). PCR reaction were performed as follows: 95°C 3 mins, 98°C 20 s, 5 cycles of 65°C 1 min and 72°C 2 min, 98°C 20 s, 5 cycles of 65°C 1 min and 72°C 2 min. The subsequent PCR product was cleaned with 1.8x AMPure XP beads (Beckman Coulter) and resuspended in 25μl of elution buffer. A second round of amplification was performed using primers targeting the 5′ sequencing adaptor (P8) and 50 bp downstream of the cleavage site (P9) for 20 cycles. The resulting PCR product was then size selected using a dual size-selection cleanup of 0.4x and 0.8x AMPure XP beads (Beckman Coulter) to remove genomic DNA and small fragments (<200 bp) respectively. This size-selected product was subsequently re-amplified to add the 3′ sequencing adaptor with primer P8 and P10 for an additional five cycles. The final PCR product was cleaned with 0.75x AMPure XP beads (Beckman Coulter) and was re-amplified to add flow-cell adaptor and sample index for 5 cycles. All PCR reactions used HiFi HotStart ReadyMix 2x (KAPA Biosystems) with the manufacturer’s recommended conditions. The library was sequenced on an Illumina NextSeq 500 sequencer using paired-end 150 cycle reads. All primers used are listed in **Table S1**. Sequence data and associated data files are deposited in Figshare with a doi link: https://doi.org/10.6084/m9.figshare.7374155,

### Sequence processing pipeline

Across three replicates, we sequenced a total of 148 million paired-end reads on an Illumina NextSeq 500. We first clustered these paired-end reads by their 15 bp UMI sequence and then filtered out reads with less than 90% identity within their representative UMI clusters. Sequence identity was identified using edlib (Šošić & Šikić 2017). UMIs with fewer than 10 reads were excluded from downstream analysis. This yielded 4,405,379 UMIs (91,325,700 reads), representing ∼61.8% of our sequencing data (**Table S2**). We then selected the most common forward and reverse read sequence for each UMI for further processing. These forward and reverse reads were merged into a single read using PEAR (Zhang et al. 2014) and aligned in a two step process as follows. First, we sought to identify the ‘reference’ sequences for each programmed array sequence. We aligned the merged reads to a backbone sequence where the guides and targets were represented by Ns using EMBO’s needleall software (Needleman & Wunsch 1970) with the following scoring matrix: match=5, mismatch=-4, gap-open=-20, gap-extension=-0.5. The mismatch penalty for Ns was set to 0. The sequence over the guide region was then extracted and matched against the list of programmed array sequences. Guide sequences with more than 2 mismatches to the designed guides were excluded, with edit distances assessed with UMI-tools (Smith et al. 2017). Second, merged reads were aligned to their discovered reference, in which Ns were replaced by the guide/target sequence identified from the first step, using Biopython.pairwise2 (Cock et al. 2009) with the following scoring matrix: match=5, mismatch=-4, gap-open=-13, gap-extension=-0.5. All indels were then right aligned (e.g. **Figure S3A**). Aligned reads with indels within −3/+2 bp of the cleavage site were assigned to their indel class. Aligned reads were excluded for downstream analysis if the sgRNA and target sequence didn’t match, the result from template switch during lentivirus transduction (Hill et al. 2018; Sack et al. 2016), or unexpected mutations introduced during synthesis, cloning, and PCR. A final library of 1.19 million unique reads (UMIs) were identified. Our library of 1,000 microhomology sequences were processed by this same pipeline, yielding a final library of 249,039 UMIs from 31,239,645 paired-end sequencing reads. Scripts and other software are available from our GitHub repository: https://github.com/shendurelab/CRISPR_NHEJ_prediction.

### Data processing and analysis

#### kpLogo Analysis

Sequence motif analysis was conducted with kpLogo (Wu & Bartel 2017) using default settings with a specified k-mer length of 1 or 2. Input sequences were weighted by the frequency of insertion.

#### Microhomology identification

For *n* from 1 to 10 nucleotides, the last *n* nucleotides upstream of each deletion were compared to the last *n* nucleotides of the deleted sequence (as the deletion is right aligned. **Figure S3A**). The length of microhomology was identified as the largest *n* nucleotides match in sequence.

### Machine learning modeling

We phrased our problem of predicting repair outcomes and their frequencies as that of a classification task with 584 classes. Because large mutation events are rare, we limited our classification effort to deletion events ≤ 30 bp, and we grouped insertions ≥ 3 bp into one class. In total, we defined 584 classes of indels. These classes include 563 deletion alleles, 4 possible single nucleotide insertion, and 16 possible dinucleotide insertion and insertions ≥ 3 bp. There are a total of 585 potential deletion events that are both ≤ 30 bp in length and overlap with the −3/+2 window around the cleavage site. We captured 563 deletion alleles in our data; the missing 22 classes are mainly large deletions. As input to our model, we defined 2,962 binary features. These are 1) 384 binary features corresponding to the one-hot encoded target sequence (excluding the PAM region), including 80 for single nucleotide content (4 nucleotides * 20 positions) and 304 for dinucleotide content (16 dinucleotides * 19 positions); 2) 2,578 binary features corresponding to MH tracts; specifically, for each of the possible deletion event class, we defined 5 binary features corresponding to the length of the MH tract, if any (0-4 bp * 563 deletion event classes, total 2,815 binary features). After excluding 237 binary features corresponding to characteristics never observed in the training data, we were left with 2,578 binary features. Our 4,611 programmed sequences were randomly partitioned into a training set of 3,750 sequences, a validation set of 450 sequences, and a test set of 411 sequences.

We trained both the logistic regression and the neural network models in a standard manner for machine learning models. However, because each target sequence can generate many possible repair outcomes, we trained our models using soft labels that correspond to the probability that each class is observed, rather than hard labels that force each input to correspond exclusively to one class. Each model was trained using the Adam optimizer (Kingma & Ba 2014) with a learning rate of 0.001 and a categorical cross-entropy loss. Training proceeded for a maximum of 100 epochs with a “patience” of 1, meaning that training was stopped after two epochs with no improvement in validation set performance. All initializations and the hyperparameters for the Adam optimizer were set to the defaults in Keras v2.0.8 (Chollet & Others 2015) with a backend of Theano v1.0.1 (The Theano Development Team et al. 2016).

We selected the best model based on performance on the validation set according to the coefficient of determination using grid search over several hyperparameters. For the logistic regression models, this search involved separate scans over regularization strengths for L1-regularization and L2-regularization individually with a range of 10^−6^ to 10^−1^(**Figure 6B**). For the neural network models, this search first involved a search over a grid of structures with between 1 and 3 layers and between 1 and 4096 nodes per layer, excluding the model with 3 layers and 4096 nodes due to memory constraints. Once the number of nodes per layer was selected, all layers in the model had that number of nodes. Each neural network model used the ReLU activation function (f(x) = max(0, x)) at the hidden layers. After the best structure was selected, a scan was performed over regularization strengths as with the logistic regression models.

## Acknowledgments

We thank members of the Shendure lab and Dezhong Deng for helpful comments and discussion. This work was supported by the Paul G. Allen Frontiers Foundation (Allen Discovery Center grant to J.S.) and grants from the NIH (R01HG009136 and UM1HG009408 to J.S. and 1K99HG010152-01 to A.M.). J.S. is an investigator of the Howard Hughes Medical Institute.

## SUPPLEMENTARY FIGURES

**Figure S1.**
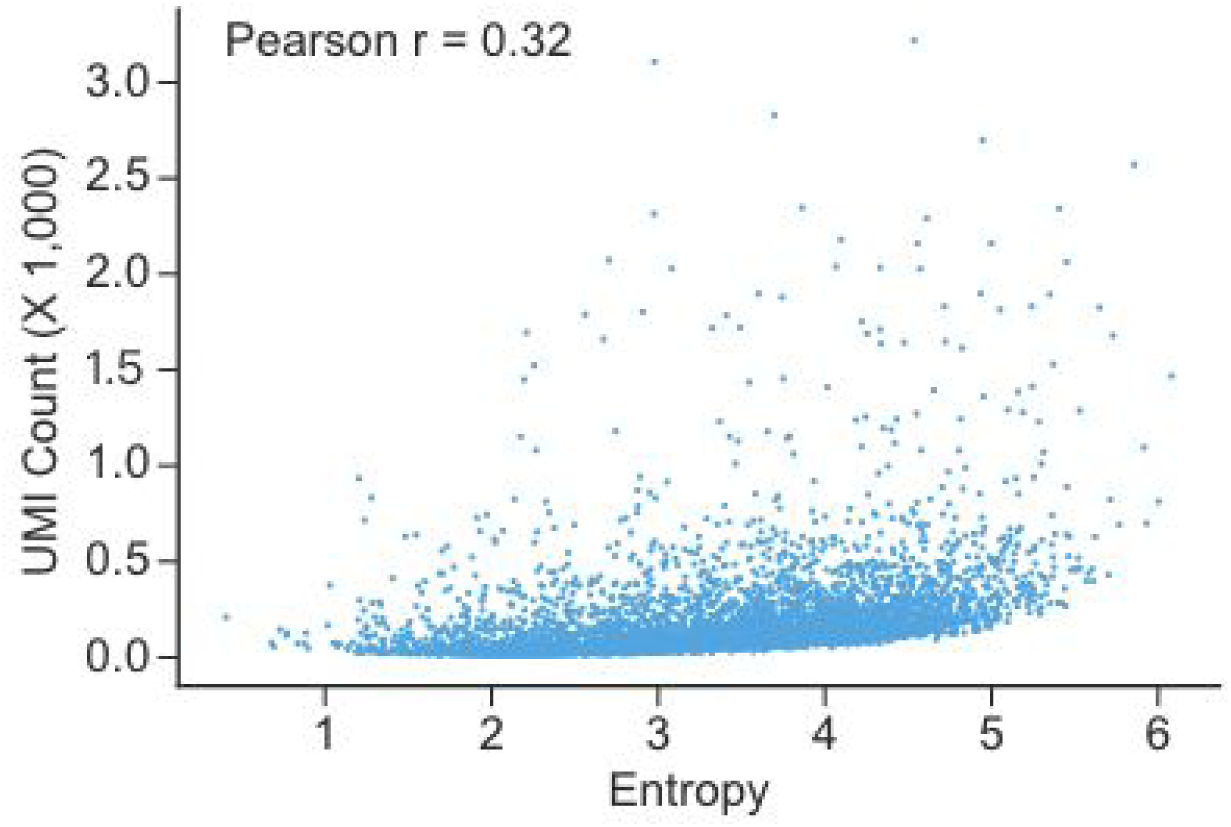
Evaluation of the correlation between entropy and UMI counts. Estimates of target-specific entropy (x-axis) are only modestly correlated with UMI counts (y-axis). Pearson r = 0.32.

**Figure S2.**
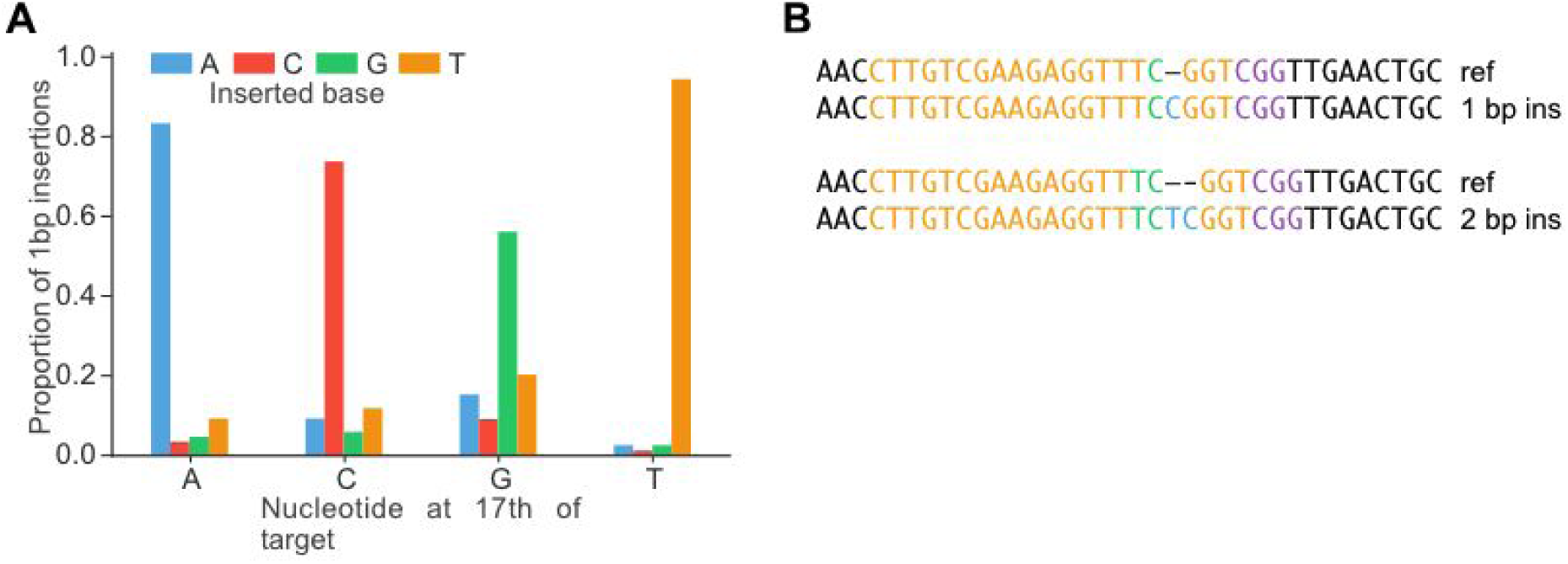
1-2 bp insertion events are templated by the nucleotides upstream of the cleavage site. **(A)** Most 1 bp insertions were predicted, and presumably templated, by the identity of the 17th nucleotide of the target sequence. **(B)** Example of insertions templated by the 17th (top) or 16th and 17th (bottom) position. Template nucleotides are shown in green and inserted nucleotides are shown in blue.

**Figure S3.**
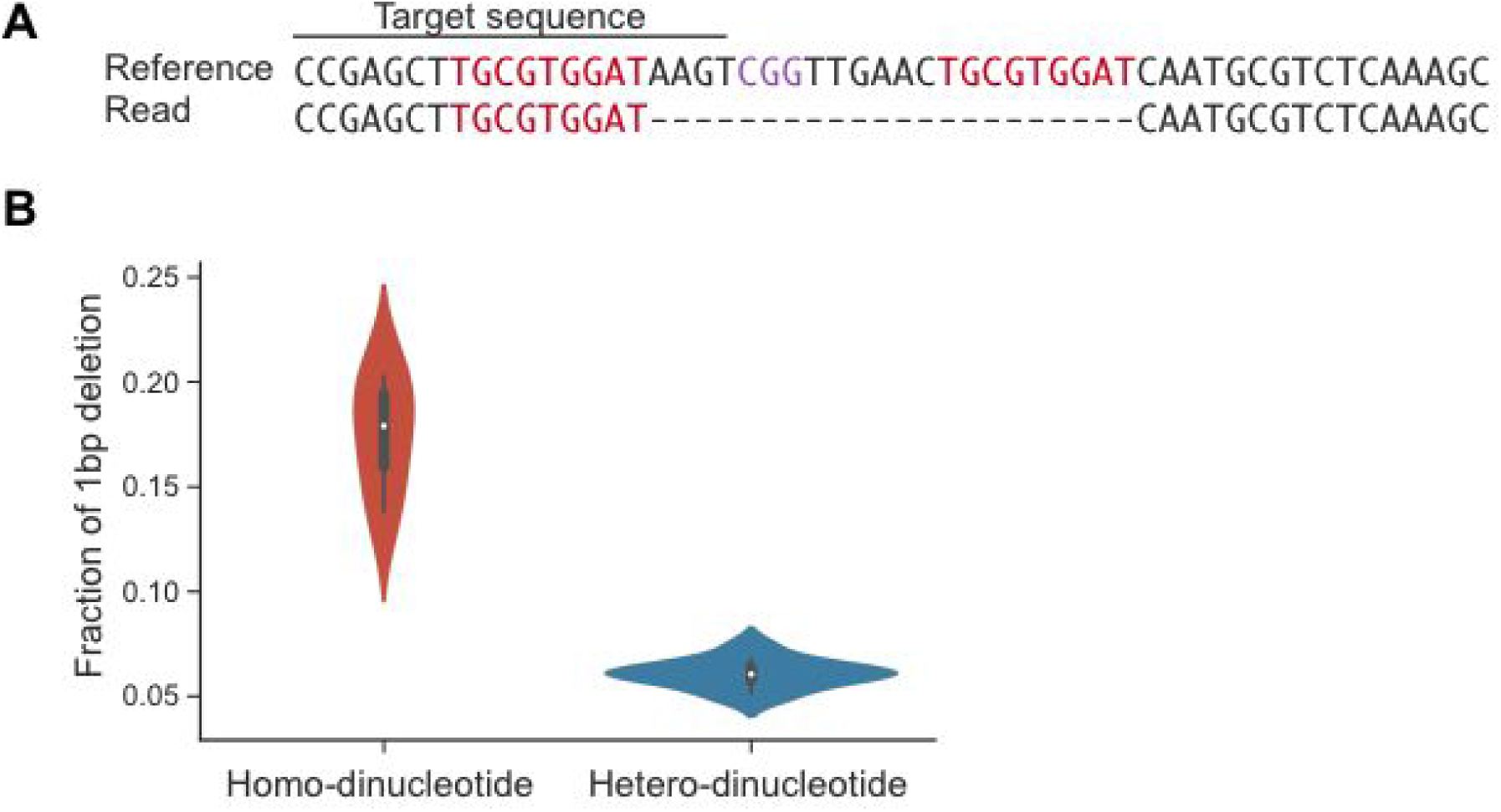
Examples of microhomology usage. **(A)** An observed example of a long MH tract mediating a deletion event. PAM and microhomology are shown in purple and red, respectively. This particular outcome, involving a 9 bp MH tract, represented 9% of indel events associated with this target. **(B)** Targets with identical nucleotides (*i.e.* ‘homo-dinucleotide’) spanning the cleavage site exhibit a much higher proportion of 1 bp deletions than targets with non-identical nucleotides (*i.e.* ‘hetero-dinucleotide’) spanning the cleavage site, suggesting that 1 bp microhomology may help mediate 1 bp deletion events.

**Figure S4.**
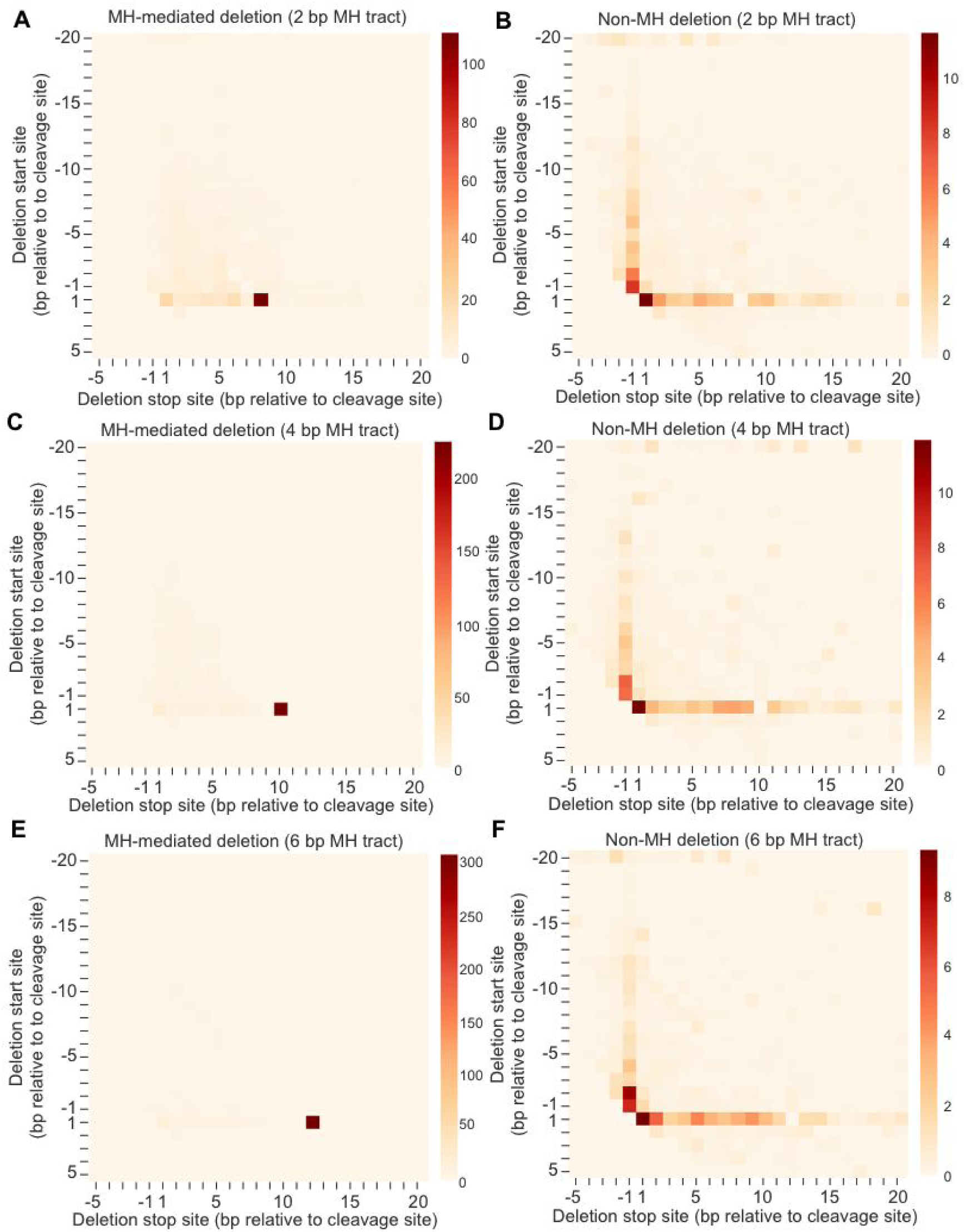
Heatmap of deletions with microhomology design. **(A,C,E)** Heatmap of showing frequency of start/stop sites of MH-mediated deletions with 2 bp, 4 bp, 6 bp programmed microhomology, respectively. **(B,D,F)** Heatmap of showing frequency of start/stop sites of non-MH deletions with 2 bp, 4 bp, 6 bp programmed microhomology, respectively.

**Figure S5.**
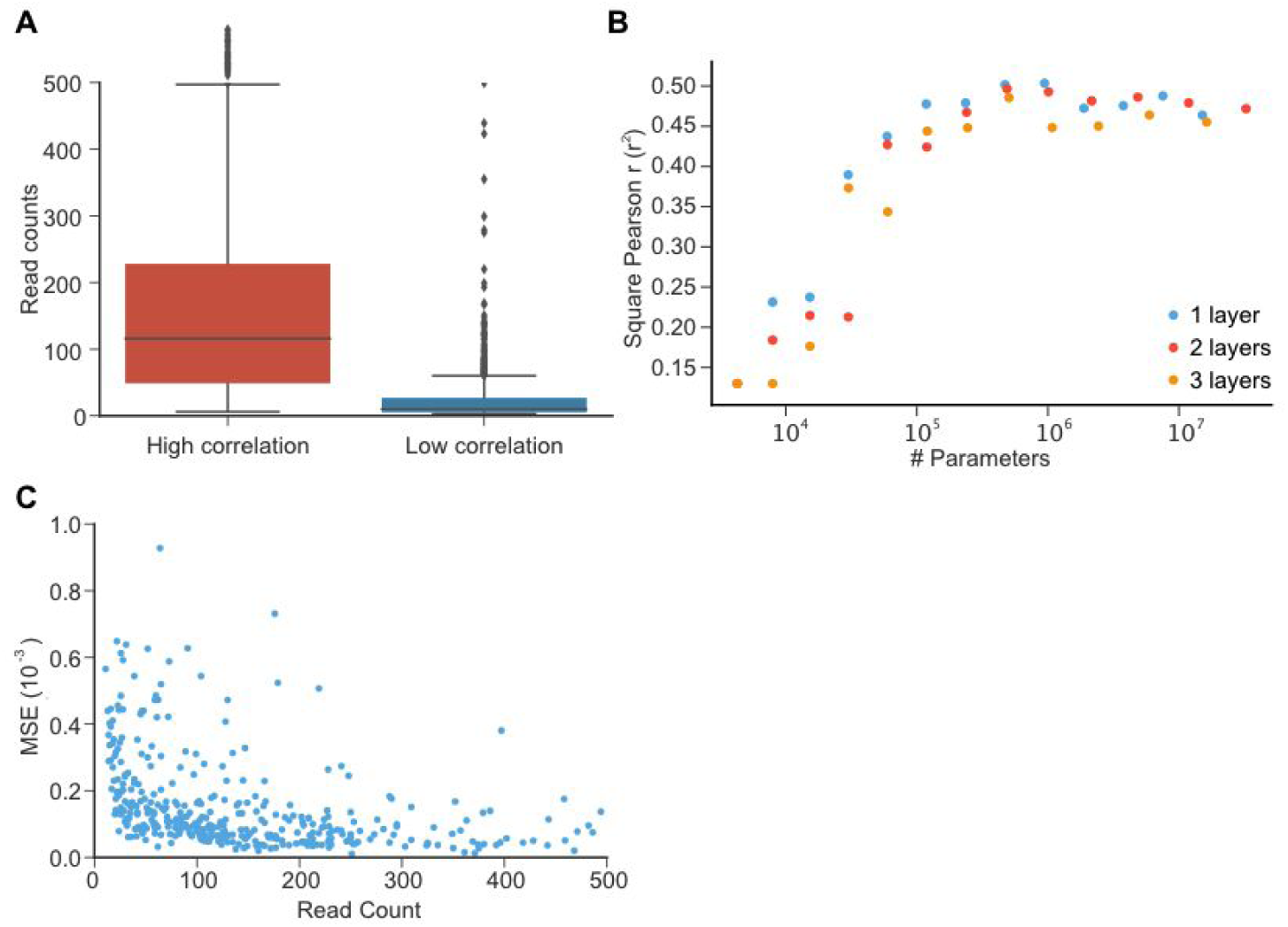
Machine learning model selection and performance. **(A)** Read counts for targets exhibiting high (r > 0.75) vs. low (r < 0.75) correlation between replicate experiments. The median read count for the two groups are 117 and 23, respectively. Targets with low correlation between replicates (r < 0.75) were excluded from model training/validation/testing. **(B)** Performance of neural network as a function of the number of parameters and layers. We used the coefficient of determination as a metric of prediction performance. **(C)** Poorly predicted targets (high MSE) largely corresponded to those that were poorly sampled.

**Figure S6.**
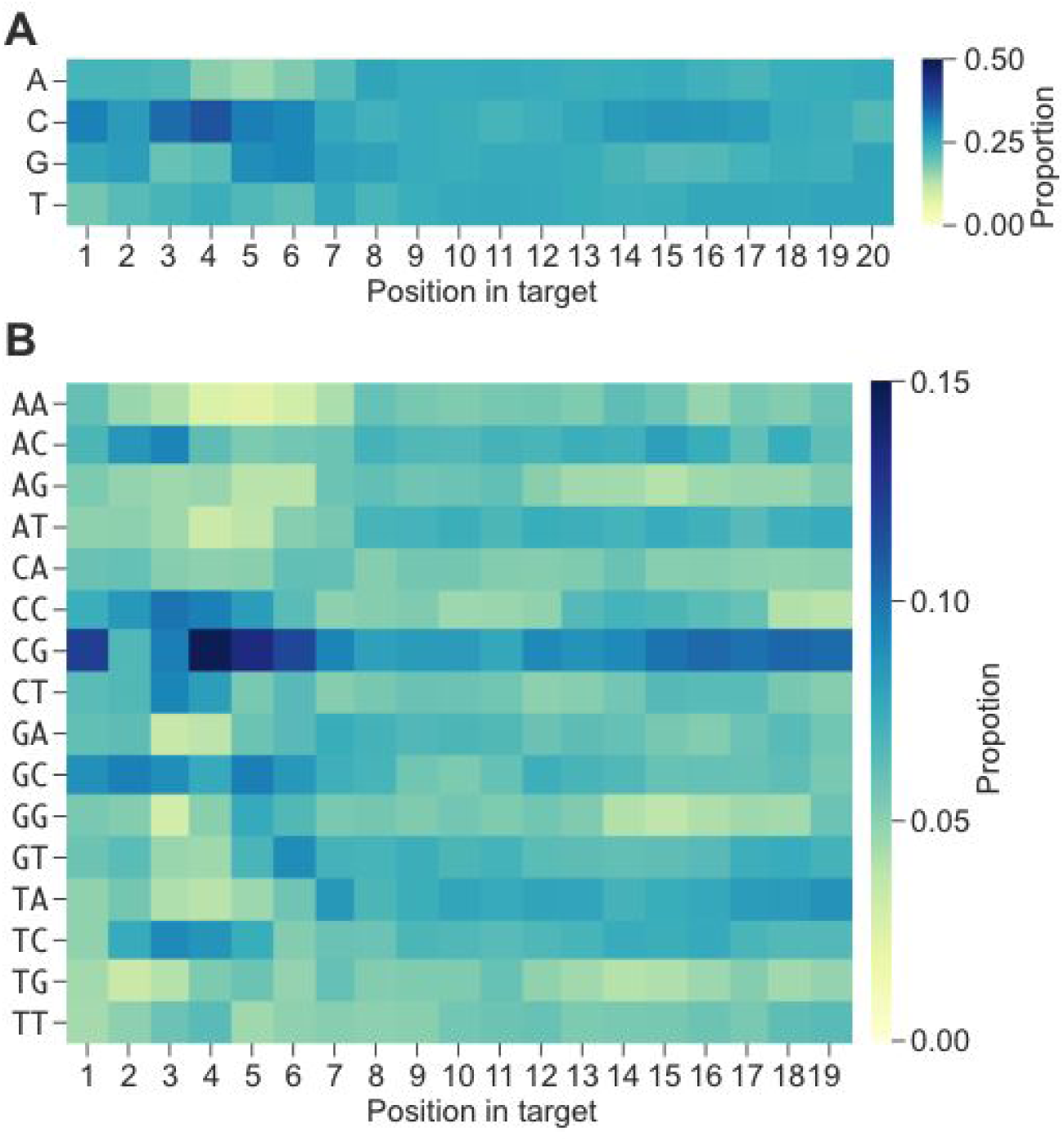
Sequence content of synthetic sgRNA-target library. **(A, B).** Heatmaps of mononucleotide (A) and dinucleotide (B) balance within the final subsampled library of 6,872 well-represented CRISPR/Cas9 targets on which most analyses were performed. Each column sums to 1. Although initially designed sequences were effectively random and balanced in mono/dinucleotide content, the overrepresentation of CG dinucleotides was likely introduced by how we screened these initial designs to remove on-target or off-target matches against the human genome (*i.e.* thereby subtly selecting in favor of designs containing CG dinucleotides, which are underrepresented in the human genome).

**Table S1.**
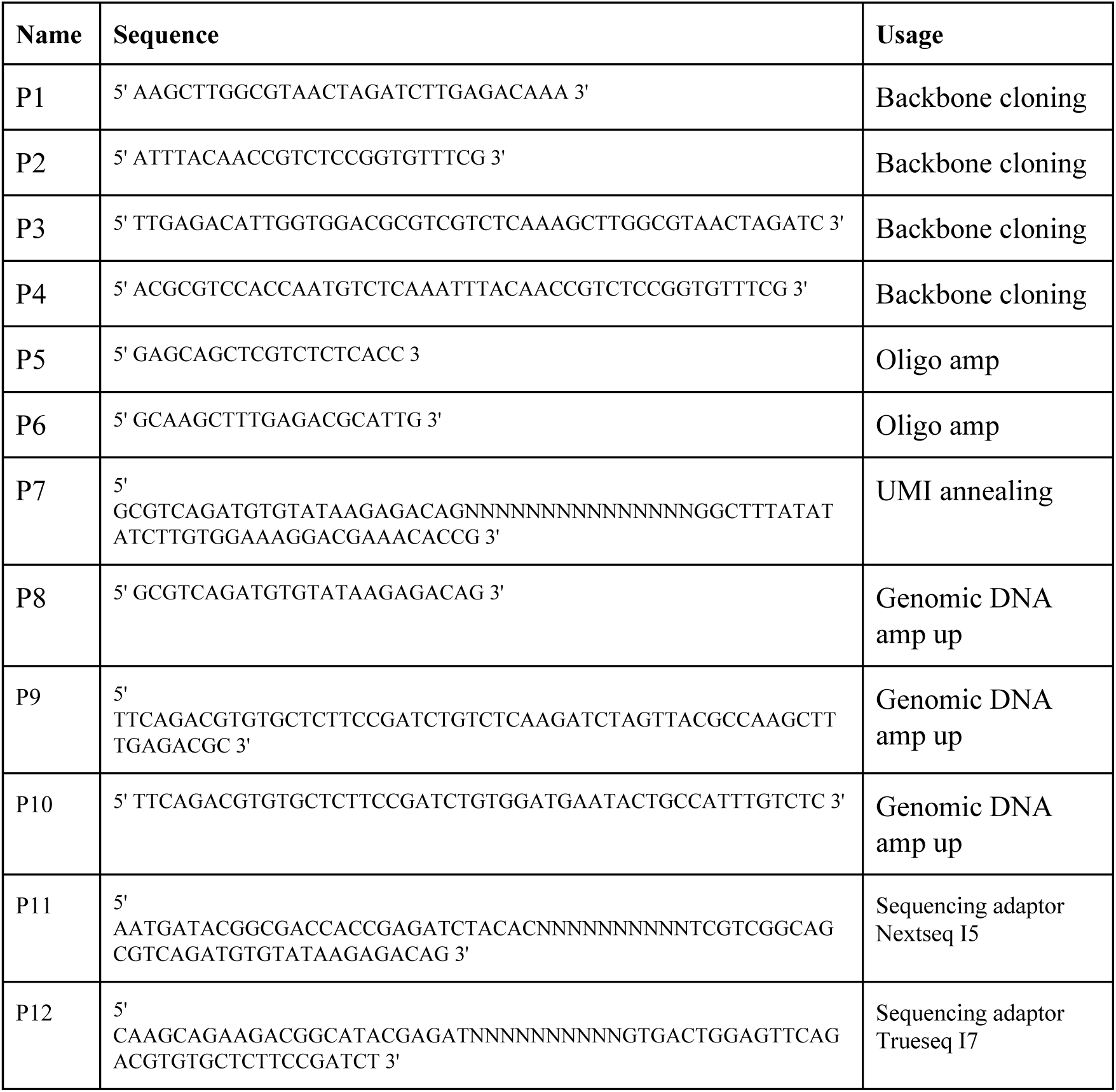
Primers. Heatmap of deletions with microhomology design.

**Table S2.** Statistics of sequencing runs.

| Sequence library | Reads | UMIs | UMI pass filter | Reads pass filter | UMI with designed sequence | Template switch | Other mutations | Final UMI count |
| --- | --- | --- | --- | --- | --- | --- | --- | --- |
| Replicate 1 | 42,796,114 | 8,393,602 | 1,327,837 | 27,896,784<br>(65%) | 1,005,606 | 27.15% | 37.28% | 357,671<br>(35.57%) |
| Replicate 2 | 60,898,224 | 12,094,298 | 1,783,500 | 39,107,335<br>(64%) | 1,354,159 | 26.98% | 37.27% | 484,056<br>(35.75%) |
| Replicate 3 | 44,075,664 | 9,794,807 | 1,294,042 | 24,321,581<br>(55%) | 982,804 | 27.92% | 36.56% | 349,066<br>(35.52%) |
| Total of Replicates 1-3 | 147,770,002 |  |  |  |  |  |  | 1,190,615 |
| MH design | 31,239,645 | 6,266,832 | 659,220 | 20,810,016<br>(67%) | 445,440 | 8.05% | 36.04% | 249,039<br>(55.91%) |

